# Identity and Nature of Neural Stem Cells in the Adult Human Subventricular Zone

**DOI:** 10.1101/2023.10.02.560459

**Authors:** Salma Baig, Javad Nadaf, Redouane Allache, Phuong U. Le, Michael Luo, Annisa Djedid, Maryam Safisamghabadi, Alex Prat, Jack Antel, Marie-Christine Guiot, Kevin Petrecca

**Author notes:** These authors have contributed equally (Salma Baig, Javad Nadaf).

## Abstract

The existence of neural stem cells (NSCs) in adult human brain neurogenic regions remains unresolved. To address this, we created a cell atlas of the adult human subventricular zone (SVZ) derived from fresh neurosurgical samples using single-cell transcriptomics. We discovered 2 adult radial glia (RG)-like populations, aRG1 and aRG2. aRG1 shared features with fetal early RG (eRG) and aRG2 were transcriptomically similar to fetal outer RG (oRG). We also captured early neuronal and oligodendrocytic NSC states. We found that the biological programs driven by their transcriptomes support their roles as early-lineage NSCs. Finally, we show that these NSCs have the potential to transition between states and along lineage trajectories. These data reveal that multipotent NSCs reside in the adult human SVZ.

## INTRODUCTION

The existence of active neurogenic sites in the human adult brain has not been resolved. Evidence for adult neurogenesis in the hippocampus (*1*) and neuroepithelium (*2*) suggest some degree of regenerative capacity.

During early human development, the SVZ emerges as a neurogenic niche containing molecularly diverse populations of RG that give rise to cells of the neocortex (*3-5*). With aging, the SVZ undergoes cytoarchitectural changes maturing into four cellular layers (*6*), and the fetal RG are thought to acquire astroglial characteristics becoming potentially adult NSCs of the SVZ (*7*).

Extensively characterized in the adult mouse brain, these SVZ NSCs, known as B cells, retain regenerative potential and can generate a variety of new-born neurons and glia (*8-10*). While the adult human brain is believed to retain NSCs of embryonic origin (*11, 12*), evidence for their existence *in situ*, particularly in the SVZ, the largest germinal region in the adult brain, is less certain. This paucity of data is primarily due to access to high-quality specimens.

To address this, we created a single-cell transcriptomic atlas of the adult human SVZ from freshly derived neurosurgical specimens. We found 4 NSC populations. Two NSC types, aRG1 and aRG2, are transcriptomically and biologically similar to fetal eRG and fetal oRG, respectively. We also discovered early neuronal and oligodendrocytic NSC states showing lineage-emergence and maturation while retaining developmental features. Biological programs underlying these states revealed neurodevelopmental, neuroinflammatory, and injury response programs. Lastly, we show that these NSCs have the potential to transition between populations and along lineage trajectories.

## RESULTS

### Adult Human Subventricular Zone Cell Atlas

SVZ samples were acquired from 15 patients undergoing brain tumor surgery. Their ages ranged from 38 to 72 years (Fig. 1A, fig. S1). SVZ regions that were normal on MRI and were to be removed during the course of surgery were sampled. The transcriptomes of 10,834 cells were acquired by single-cell RNA-sequencing (scRNA-seq). Cells containing cancer type-specific copy number aberrations were removed from further analysis (fig. S2).

**Figure 1.**
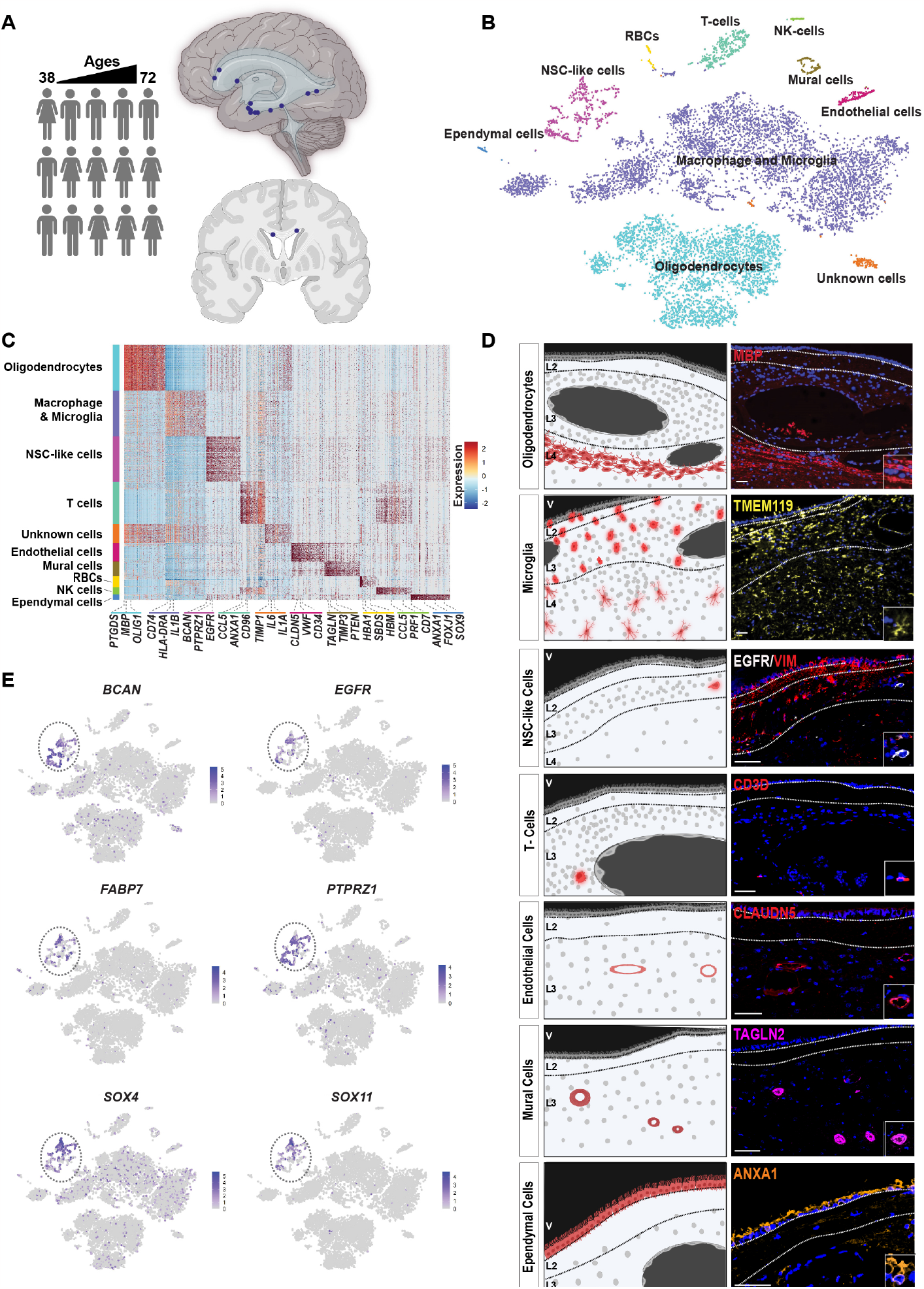
Adult Human Subventricular Zone Cell Atlas. **A**. Schematic diagram illustrating source and brain regions of adult human SVZ neurosurgical samples. **B**. I-distributed Stochastic Neighbor Embedding visualization (tSNE) of adult SVZ cells colored by cell types. NSC - neural stem cell, NK - natural killer, RBC - red blood cell. **C**. Heatmap of top 100 genes expressed in each SVZ cell type. **D**. Localization of cell types within normotypic autopsy SVZs using cell-type specific protein markers. Scale bar = 50µm. **E**. Expression of indicated genes in all SVZ cell type clusters visualized on tSNE.

The Louvain algorithm resolved 10 clusters (Fig. 1B). Cell-types were labelled based on expression of canonical gene markers: oligodendrocytes; macrophage/microglia; neural stem cell (NSC)-like; T cells; endothelial cells; mural cells; red blood cells; natural killer cells; ependymal cells; and an unidentified cell cluster (Fig. 1C, table S1). Cell-type specific protein markers were used to spatially locate these cell types in human adult SVZ samples derived from normotypic autopsy brain tissue (Fig. 1D). Consistent with the known cellular composition of the SVZ (*6*), oligodendrocytes were most abundant in layers 4 and 3, respectively. NSC-like cells were detected in the astrocytic band, layer 3. The astrocytic band in the human adult SVZ has been reported to contain NSCs (*7*).

Thirteen of the 15 patients contributed cells to the NSC-like cluster. Cells in this cluster expressed canonical neural progenitor and RG genes such as *BCAN* (64% of cells), *SOX4* (68% of cells), *SOX11* (47% of cells), *PTPRZ1* (69% of cells), *FABP7* (43% of cells), and *EGFR* (43% of cells) (Fig. 1E) (*3-5, 13*). However, no gene was expressed in all cells in the NSC-like cluster, suggesting cellular diversity within this cell cluster.

### Neural Stem Cell Diversity Within the Adult Subventricular Zone

Since differentiated cells in the SVZ cell dataset forced diverse NSC-like cells into one cluster, we created a progenitor-only dataset using the adult NSC-like cluster and enriched it with fetal progenitor/stem cells from Nowakowski *et al*. (2017) and Couturier *et al*. (2020) (*4, 9*) (Fig. 2A). In this merged dataset the adult NSC-like cluster resolved into 4 clusters (C4, C5, C7, C9) (Fig. 2B). C5 contained cells from the adult SVZ and fetal eRG, and C9 contained cells from the adult SVZ and fetal oligodendrocyte progenitor cells (OPCs). To preserve the pure adult gene signatures and account for the transcriptomic differences between adult and fetal cells, all fetal cells were removed from further downstream analysis of the 4 SVZ NSC-like cell clusters.

**Figure 2.**
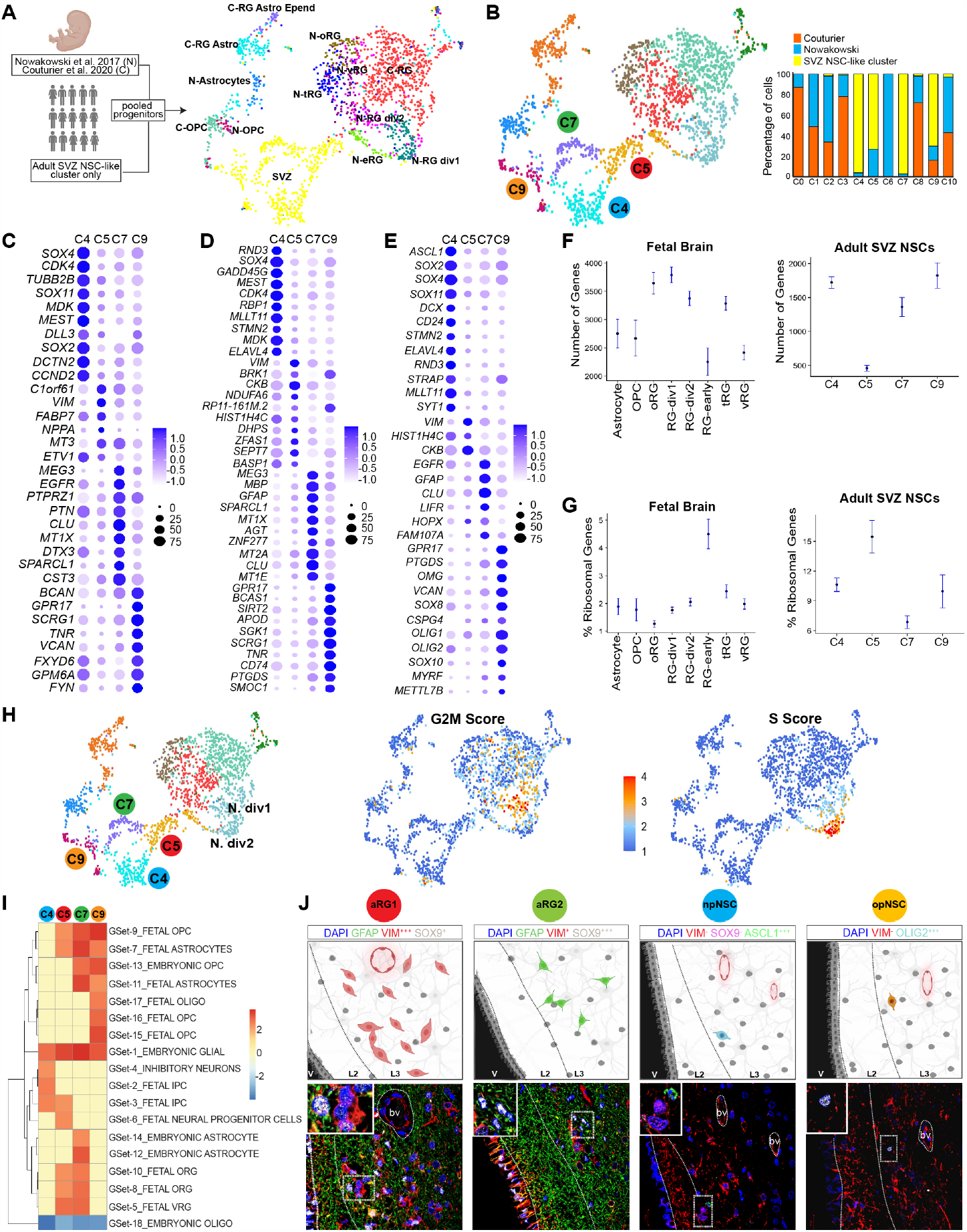
Neural Stem Cell Diversity Within the Adult Subventricular Zone. **A**. Schematic diagram illustrating sources of samples from adult human SVZ regions and fetal brains, and Uniform Manifold Approximation and Projection (UMAP) visualization of the merged progenitor cells selected from the adult SVZ NSC-like cluster and the fetal datasets colored by original cell annotations. **B**. UMAP visualization of the merged progenitor cells selected from the adult SVZ NSC-like cluster and fetal datasets by clusters. Bar graph shows the dataset source of the cells contributing to each cluster. **C**. Dot plot of the top 10 genes expressed in each of the 4 NSC clusters compared to all SVZ clusters. **D**. Dot plot of the top 10 genes expressed in each of the 4 NSC clusters compared to NSC clusters. Ribosomal genes are not shown (see methods). Redundant genes are shown only once. **E**. Expression of curated known gene markers of RG, NSCs, neuronal progenitors and oligodendrocyte progenitors. **F**. Transcriptome size of adult NSC-like and fetal brain cells. Confidence interval = 95%. **G**. Ribosomal gene content of adult NSC-like and fetal brain cells. Confidence interval = 95%. **H**. Cell cycle phase scores calculated based on expression of G2M and S markers. **I**. Summary of gene set enrichment analysis using curated human brain cell types published single-cell RNAseq datasets. Dendrogram shows unsupervised hierarchical clustering of the data. Color bar indicates normalized enrichment score. Red and blue colors represent significant enrichment or down-regulation (FDR<0.0001), respectively, yellow indicates no significant result. **J**. Localization of aRG1, aRG2, npNSCs, and opNSCs within normotypic autopsy SVZs using cell type-specific protein markers. Scale bar = 50µm.

Differential gene expression analysis was then used to determine NSC cluster-specific expression patterns (Fig. 2C, table S2). The most highly differentially expressed genes in each of the 4 adult NSC clusters, compared to all other SVZ cells, were: C5 - *C1orf61, VIM, FABP7, NPPA, MT3*; C7 - *MEG3, EGFR, PTPRZ1, PTN, CLU*; C4 - *SOX4, CDK4, TUBB2B, SOX11, MDK*; C9 - *BCAN, GPR17, SCRG1, TNR, VCAN*.

### Transcriptomic Differences between NSC Clusters Show Radial Glial Identity and Lineage-Specific Programs

To explore the transcriptomic differences between the NSC clusters, we identified genes that were differentially expressed between these clusters. C5 showed high expression of classical RG markers *VIM* and CKB (Fig. 2C, D and E; table S3) (*3-5*). Ribosomal genes *RPS3A, RPLP1, RPS14, RPL23A, RPS27A*, and *RPL39* were also highly expressed, as were the mouse radial precursors ribosomal genes *RPL31, RPS15* and *RPL3* (*15*). No lineage-specific genes were expressed. Since C5 contained adult SVZ NSCs and fetal eRG from gestational week 8 (*4*), we compared transcriptomic features of these cells to other SVZ and fetal cell types. We found that both adult and fetal C5 cells shared low transcriptome sizes (Fig. 2F) and high expression of ribosomal genes (Fig. 2G). The uncoupling of protein synthesis and ribosome biogenesis has been reported in activated and quiescent stem cells, albeit in opposite directions (*14*). Likewise, RG precursors from mouse embryos at embryonic week E11.5 have been shown to express higher levels of ribosomal biogenesis genes (*15*). Some cells in C5 expressed a cell cycle gene signature (Fig. 2H).

The C7 transcriptome contained classical astroglial marker genes expressed in human embryonic and murine adult NSCs such as *ALDOC, ID3, GFAP*, and *GLAST* (*15-18*). Fetal oRG genes *HES1, HOPX, LIFR, SLC1A3, HEPACAM, FAM107A*, and *TNC* were also highly and uniquely expressed in C7 (Fig. 2D and E; table S3) (*4, 5*). Adult SVZ NSCs have been reported to retain astroglial characteristics (*16, 19*). C7 also expressed extracellular matrix-related genes PTN, *BCAN, NCAN*, and *SPARCL1* (*20*), and gap/tight junction genes *CLDN1/12, COX43/GJA1*, and *TJP1* (table S3) (3, 21, 22).

C4 expressed genes associated with embryonic development and neuronal programs such as *ASCL1, SOX2, CD24, SOX4*, and *SOX11* (Fig. 2C, D and E; table D3) (*23-25*), and neurogenic oRG transcription factors *HES6, NHLH1*, and *CBFA2T2* (table 3) (26). C4 also expressed *BEST3, STMN2, ELAVL4*, and *DCX*, gene markers of fetal intermediate neuronal progenitors/new-born neurons, and *ZIC2, MOXD1*, and *C1orf61*, markers of ventral telencephalic RG (*4*). These cells did not express classical fetal newborn interneuron markers *PAX6, TBR2*, or *SATB2*, suggesting differences between fetal and adult-born neurons. *RND3, MLLT11, STRAP*, and *SYT1*, markers of neuronal trajectory maturation critical in cortical evolution for neurite outgrowth and axonogenesis, were differentially highly expressed in C4 (*23, 27-32*), whereas expression of RG/neural progenitor markers *VIM, GFAP*, and *HES1* were downregulated. Markers of adult human neurons *CALB2, CCK, PVALB, SST*, and *VIP* were also absent (*33*). C4 also expressed markers of cell cycle activity including *CCND1, CCND2, TOP3A*, and *CDK4*, and some cells expressed a cell cycle gene signature (Fig. 2H).

C9 cells highly expressed OPC markers *BCAN, CSPG4, OLIG1/2*, and *SOX10* (Fig 2C, D and E; table S3) (*34-36*). OPC, preOPC, and differentiation-committed OPC markers *VCAN, SOX6, PCDH15, MEGF1*, and *GPR17* were uniquely expressed in C9, as were migration-related genes *TNS3* and *FYN* (*33, 37, 38*). Premyelinating oligodendrocyte genes *BCAS1, SGK1, TCF7L2*, and *SCRG1* were also expressed in C9 (*39, 40*), as were genes involved in neuro-immune processes such as *CD74, HLA-DRB1*, and *CXCR4* (*41*). Demyelination has been shown to cause adult murine OPCs to express immune cues contributing to the post-injury inflammatory milieu and to support migration (*42*).

### Adult Neural Stem Cell Gene Signature Comparison Using Gene Set Enrichment Analysis

We next compared gene signatures of the 4 NSC clusters to curated genesets of human embryonic and fetal brain cell types sourced from published scRNA-seq studies (*4, 43-45*). All 4 NSC clusters showed enrichment for a NSC-like signature and low expression for differentiated oligodendrocytes (Fig. 2I). C4 showed enrichment for 3 neuronal progenitor programs. C9 showed enrichment for oligodendrocyte progenitor programs. C7 most strongly enriched for embryonic astrocytes and fetal RGs. C5 was enriched for RG programs and a neural progenitor program.

Based on these gene expression patterns and comparisons to available datasets, we labelled C5 cells as adult radial glia 1 (aRG1) and C7 cells as aRG2. C4 and C9 cells have acquired lineage specificity and because of this potential transient nature we labelled these as progenitor states, C4 as neuronal progenitor NSCs (npNSCs) and C9 as oligodendrocytic progenitor NSCs (opNSCs).

Using the differentially expressed genes as a guide, we selected protein markers to locate these 4 NSC populations within the SVZ from normotypic autopsy adult human brain samples (Fig. 2J). aRG1 (VIM^+++^, SOX9^+^) and aRG2(VIM^+^, SOX9^+++^) were distributed in layer 3 (7). opNSCs (OLIG2^+++^, VIM^−^) and npNSCs (ASCL1^+++^, VIM^−^, SOX9^−^) were found in layers 3 and 2, respectively.

### Uncovering Neural Stem Cell Programs Using Biological Program Analysis

We also characterized the biological programs inherent in aRG1, aRG2, opNSCs, and npNSCs using their most differentially expressed genes (Fig. 3; table S4). npNSCs showed enrichment for programs such as forebrain development, cerebellar cortex development, transcriptional regulation by RUNX1 and 3, pre-NOTCH expression and processing, hedgehog ligand biogenesis, and NOTCH signaling. Cell cycle processes were also active, consistent with their cell cycle score (Fig. 2H), as were neuronal processes including dendrite development, neuroepithelial cell differentiation, cell fate determination, regulation of neuron differentiation, embryonic morphogenesis, cadherin binding, and cell fate specification. These programs have been reported to be required for neuronal differentiation and maturation (*27, 29, 30*).

**Figure 3.**
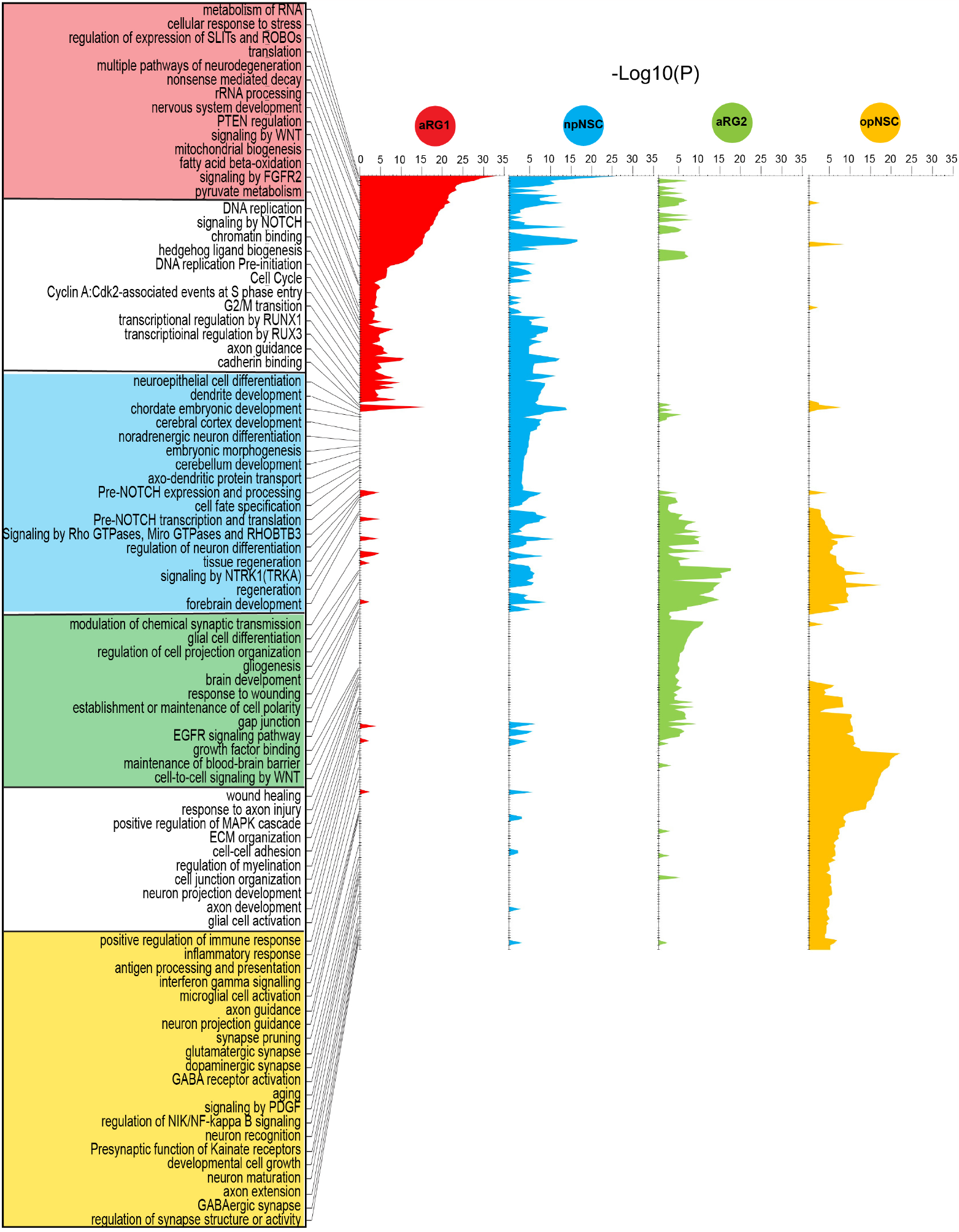
Enriched Neural Stem Cell Biological Programs. Biological program analysis performed for aRG1, npNSCs, aRG2, and opNSCs, using a maximum of the top 500 differentially expressed genes. Unique pathways/process are boxed and colored according to their state. Shared pathways/processes are boxed white.

opNSCs were enriched for processes such as gliogenesis, regulation of myelination, extracellular matrix organization, PDGF signaling, GABA B receptor activation, presynaptic function of kainate receptor, neuron recognition and maturation, axon development, synapse organization, and response to axon injury (Fig. 3). While embryonic OPCs lack ion channels or glutamatergic receptors, these channels have been shown to emerge in OPCs postnatally (46). opNSCs also expressed neuroimmune processes including glial cell activation, cytokine signaling, antigen processing and presentation, inflammatory response, and microglial activation. OPCs have been shown to modulate neuroinflammation, immune response, and central nervous system repair (47, 48). Cell motility processes, regulation of locomotion and cell motility, RAC1 GTPase cycle, and cell junction disassembly were enriched in npNSCs and opNSCs.

Although aRG1 and aRG2 shared astroglial markers, their biological programs differed (Fig. 3; table S4). aRG1 were enriched for rRNA processing, mRNA splicing, translation, ribosome biogenesis, and nonsense mediated decay. Ribosome biogenesis has been shown to be crucial for controlling stem cell homeostasis. In adult stem cells the level of ribosome biogenesis and protein synthesis determines the switch between quiescent and activated states with self-renewal decisions (*14, 49, 50*), while senescence is associated with shutdown of ribosome biogenesis (*14, 50*). In addition, active metabolic processes such as oxidative phosphorylation, mitochondrial biogenesis, pyruvate metabolism, fatty acid beta-oxidation, and stress-related processes were also enriched in aRG1. Quiescent neural stem cells maintain high levels of mitochondrial fatty acid β-oxidation and express proteins involved in mitochondrial metabolism (*49, 51, 52*). Mitochondrial dynamics have been reported to affect self-renewal and fate choices (*53, 54*). aRG1 also showed enrichment for cell cycle, consistent with their cell cycle score (Fig. 2H), and DNA replication, chromatin binding, and axon guidance, albeit to a lesser degree than npNSCs.

aRG2 were significantly enriched for brain development, EGFR signaling, response to metal ions, gap junction, cell polarity, transepithelial transport, cell-cell adhesion, and maintenance and transport across the blood-brain barrier (Fig. 3; table S4) (*55-59*). SVZ astrocytic NSCs have been reported to regulate the niche (*60*). Developmental processes such as cell-cell signaling by WNT, nervous system development, signaling by NTRK1, regulation of neurogenesis, regulation of gliogenesis and glial cell differentiation, neuron projection development, kinase activity and regulation of growth, and wound healing and tissue regeneration processes were also enriched (*56, 58, 61-64*).

### Subventricular Zone Cell to Cell Interactions

Cell to cell interaction analysis revealed interactions involving receptors expressed on NSCs known to be important in neurodevelopment, including EGFR, NOTCH, PDGFRA and PTPRS (Fig. 4, table S5). Corresponding ligands were expressed by NSCs and other niche cells. Consistent with the transcriptomic analyses, aRG1 showed relatively fewer interactions compared to aRG2, npNSCs, and opNSCs.

**Figure 4.**
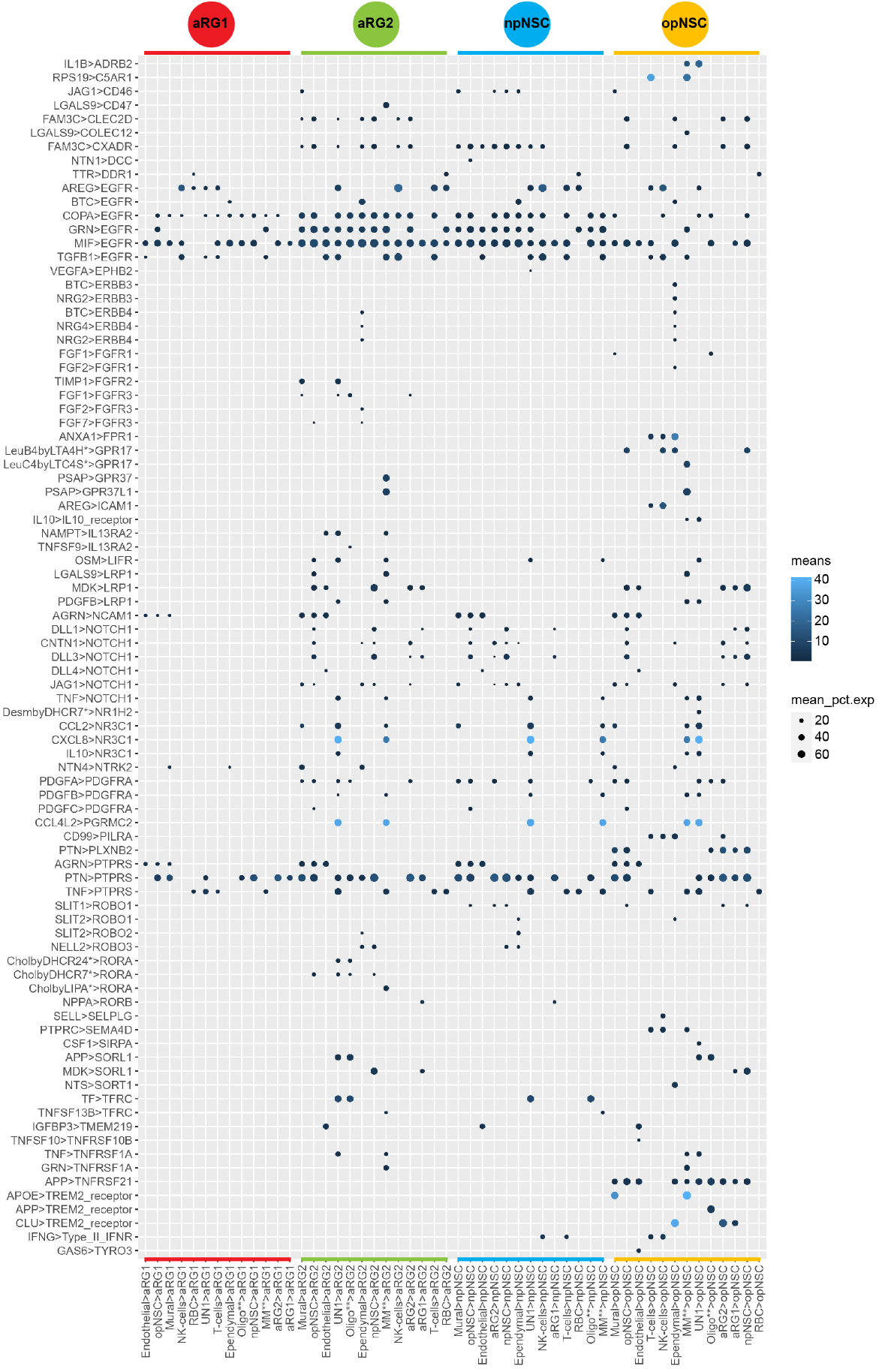
Subventricular Zone Cellular Interactions. Dot plot showing the ligand-receptor interactions from all niche cell types to NSCs. Scale bar is the mean expression of the ligand-receptor pair. Dot size is the mean of percentage of expressed cells for each cell type pair. MM^**^ - Macrophage and Microglia, Oligo^**^ - Oligodendrocyte, UN1 - unidentified.

EFGR expression was common across NSCs, but most abundant in aRG2 and npNSCs. Six ligands interacted with EFGR. Betacellulin (BTC), expressed by ependymal cells, has been reported to be involved in regulating various processes ranging from reproduction to the control of NSCs (65). Amphiregulin (AREG), expressed mainly by NK-cells and T-cells, is a mitogen for astrocytes and has been shown to be involved in embryonic morphogenesis and wound healing by interacting with EGFR (66). Progranulin (GRN) is associated with processes including embryogenesis, tumorigenesis, inflammation, wound repair, and neurodegeneration (*67, 68*).The Macrophage Migration Inhibitory Factor (MIF) interacted with all NSCs. Although its activity has not been studied extensively, it has been reported to block EGFR activation (*69*). The actions of TGFβ1 and COPA on EGFR are also not well understood.

Pleiotrophin (PTN), known to be associated with new-born neuron development (*70*), was expressed by many niches cell types and interacted with the Protein Tyrosine-Phosphatase Receptor S (PTPRS) expressed on all NSC types. PTPRS has been shown to be involved in axonogenesis, axon guidance, and adult nerve repair (*71, 72*).

Interactions exclusive to opNSCs were apolipoprotein E (APOE) from mural cells and macrophage/microglia, and clusterin (CLU) from ependymal cells, interacting with the Triggering Receptor Expressed on Myeloid cells 2 (TREM2), implicating opNSCs in response to injury, neuroprotection, and degenerative pathologies (*73, 74*).

### Neural Stem Cell Transition Capacity

We used RNA velocity to investigate trajectories between SVZ NSCs and their progeny. To reconstruct the spectrum of neuronal cell differentiation, we created a merged scRNA-seq dataset containing adult SVZ cells, fetal brain cells (*9*), and adult brain astrocytes (*75*) and oligodendrocytes (*75*) along the spectrum of differentiation. Three groups of state transition patterns were found (Fig. 5A, 5B). Group 1 cells show multistate transition capacity. All SVZ-derived NSCs were in group 1. We found that aRG1 can transition to fetal intermediate neuronal progenitor cells (IPCs), npNSCs, fetal RG, fetal OPCs, and fetal astrocyte progenitor cells (APCs). aRG2 can transition to npNSCs, fetal APCs, fetal OPCs, and adult astrocytes. opNSCs showed transition capacity to npNSCs and fetal OPCs, and adult early oligodendrocytes to a lesser degree. npNSCs showed weak transition capacity to fetal IPCs and fetal OPCs. Group 2 cells show only intra-lineage transition capacity. For example, fetal IPCs can give rise to newborn excitatory neurons. Group 3 cells show no transition capacity (Fig. 5A). For example, oligodendrocyte cluster 6 derived from the adult SVZ, a fully differentiated cell type, did not give rise to any other cell type in this dataset.

**Figure 5.**
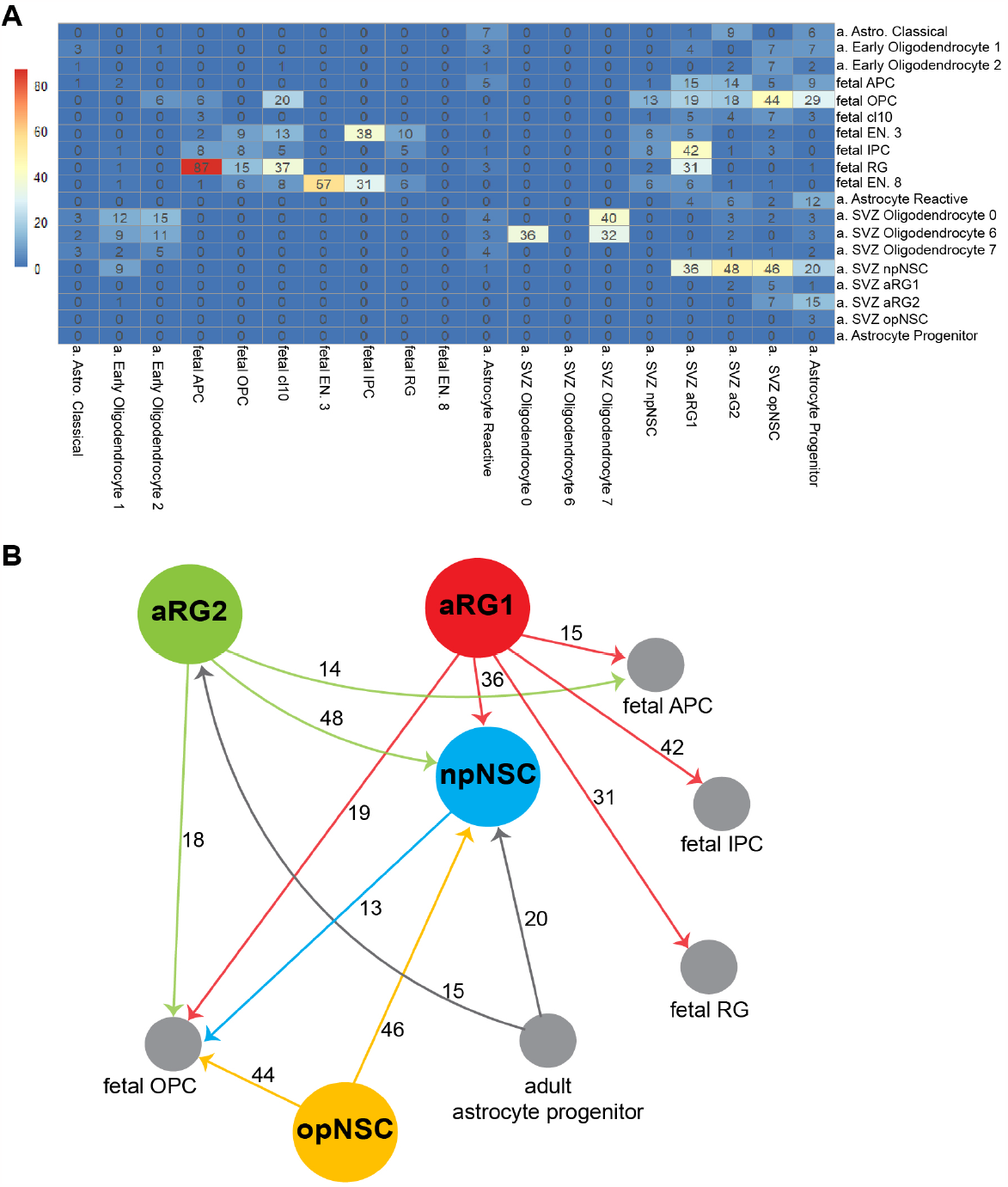
Neural Stem Cell Transition Capacity. **A**. RNA velocity analysis for adult and fetal brain cell types. Numbers are transition confidence ranging from Oto 100. **B**. Summary schematic showing cell transitions with confidences greater than 10. a - adult, astro - astrocyte. APC - astrocyte progenitor cell, OPC - oligodendrocyte progenitor cell, **EN** - excitatory neuron, **IPC** - intermediate progenitor cell, **RG** - radial glia, **SVZ** – subventricular zone.

## DISCUSSION

We have provided a cell atlas of the adult human brain SVZ. We have identified 4 NSC populations using transcriptomic and biological program analyses. aRG1, like fetal eRG, showed hallmarks of dormancy and localized to the SVZ astrocytic band (Layer 3). aRG2 were similar to fetal oRG, showing features consistent with mouse NSC populations (*8-10*). npNSCs and opNSCs showed early lineage-specification toward neuronal and oligodendroglial differentiation, respectively. We have also provided a trajectory analysis that demonstrates the intercellular and differentiation transition capacity of these NSC populations.

These data shed light on the potential differences between developmental and regenerative programs in adult SVZ NSCs, and uncover the post-developmental potential of the adult human brain. These data have important implications for our understanding of neurodegenerative, neuroinflammatory, neuroimmune, and neurooncological processes (*75, 76*).

Importantly, although the cells captured in this study were isolated from SVZ regions that are normal on MRI, they may have been impacted by their proximity to brain tumors. It is possible that signals derived from the tumors may be relevant. If true, these signals may reveal the potential of NSCs to become activated, divide, and differentiate in the adult brain.

## Supplementary Materials

### Materials and Methods

#### Adult Human Subventricular Zone Sample Collection and Processing

All samples were obtained from surgeries performed at the Montreal Neurological Institute-Hospital under a research ethics board approved protocol NEU-10-066. Consent was given by all patients. Preoperative magnetic resonance imaging (MRI) was performed for surgical planning. SVZ regions that were normal on MRI and were to be removed during the brain tumor resection surgery were sampled.

For each sample, an aliquot of cells was assessed for viability with calcein-AM and ethidium-homodimer1 (P/N L3224 ThermoFisher Scientific). Single-cell capture was performed following the Single Cell 31⍰ Reagent Kits v2 User Guide (CG0052 10X Genomics). Cell barcodes and unique molecular identifiers (UMI) barcodes were demultiplexed and single-end reads aligned to the reference genome, GRCh38, using the CellRanger pipeline (10X Genomics).

#### Single-Cell RNA Sequencing Data Processing

We applied similar approaches for adult SVZ (S, 10 834 cells), fetal (F, 12544 cells) and merged (M, 1965 cells) datasets. For each cell, counts (number of UMIs) per gene were normalized to the total counts of the cell and scaled by 10000. The data were natural-log transformed (log(counts+1)). All cells expressing more than 200 and less than 3500 genes, with lower than 8% expression for mitochondrial genes were included for further analyses. The most significant PCAs were selected (S:35;F:30;M:20), using jackstraw method and elbow visualization to cluster the cells based on a shared nearest neighbor (SNN), applying the Louvain algorithm as described in (*77*) and implemented in (*78*), with the resolution of 0.6-0.8 (S:0.6;F:0.6;M:0.8). Markers, for each cluster/program compared to all other programs, were identified using the FindAllMarkers function with default Wilcoxon test (*78*). We also applied the FindAllMarkers function within the 4 NSC clusters to highlight the difference between the 4 clusters. Cell types annotations were assigned to each cluster using known markers shown in the Figure 1C, which were also consistent with markers previously reported (*4*). For visualization purposes, t-distributed stochastic neighbor embedding (t-SNE), and Uniform Manifold Approximation and Projection (UMAP) were used. Regarding the M dataset, since the single-cell platforms were different (Fluidigm C1 vs. 10x datasets), we regress out the effect of platforms (Seurat: vars.to.regress=platform). Regarding the velocity dataset, where we added progenitor cells from a non-cancer adult brain dataset (n=322), we did not perform clustering and we only used the previous cell type annotations for each cell (n total=4593; SVZ:940, fetal:3331).

#### Copy Number Alteration Analysis

InferCNV (V.1.4.0) (*79*) was used to detect large-scale chromosomal copy number alterations (CNAs) at single-cell level using gene expression data. Gene expression values were smoothed on a moving window of 101 genes and CNAs were analyzed relative to the reference cells. The three largest clusters, Oligodendrocyte, Macrophage/Microglia, were used as reference cells. The most variable chromosomes, those with highest number of alterations, were chromosomes 7 and 10, consistent with glioblastoma. Cells with any chromosomal alterations on chromosome 7 (gain) and chromosome 10 (loss) were defined as pathological cells and were excluded from further analyses.

#### Cell Cycle Analysis

We used gene signatures for G2M and S phase, as reported by Tirosh et. al. (*80*), to quantify cell cycle phase score. Cells scoring low for both phases were considered to be in G1/G0 phase. We used CellCycleScoring function of Seurat with default parameters.

#### Ribosomal Gene Quantification

The proportion of ribosomal protein genes (Ribosomal Protein Large/Small subunit) were calculated using Seurat (PercentageFeatureSet (pattern = “RPS|RPL”)). Since the datasets were generated using different single-cell platforms, the analyses were performed separately for each dataset.

#### Gene Set Enrichment Analysis

Differentially expression genes were identified for each of the 4 NSC clusters compared to all other SVZ cells using the FindAllMarkers function with default Wilcoxon test. The genes were ranked based on average fold change followed by gene-set enrichment analysis (GSEA) (*81*). We used curated gene sets of known human brain cell types from single-cell RNA-Seq studies as reported in Molecular Signatures Database (MsigDB,C8) (*82*), as well as all signatures in fetal brains reported by Nowakowski et al. (*4*). We used GSEA-Preranked (V4.1.0; Build 27) with default parameters and 10000 permutations.

#### Cell-Cell Interaction Analysis

We used CellPhoneDB (Garcia-Alonso et al., 2021), a repository of curated receptors, ligands, and their interactions, to infer cell-cell communication networks between cell types. We provided CellPhoneDB v3.1.0 the RC-normalized count matrix by Seurat using genes with ENSEMBL ID along with cell type identities and ran the statistical analysis with the following parameters: –iteractions 10000 –result-precision 10 –threads 10 –threshold 0.1. We selected interactions with p-value < 0.05, and selected ligand/receptor interactions. Only interactions to NSCs were considered.

#### RNA Velocity Analysis

We performed velocity analysis using partition-based graph abstraction (*83*). In the first step, reads were aligned to the reference genome, GRCh38, using the CellRanger pipeline form 10X Genomics as previously described (*75*). Spliced and un-spliced counts were calculated and saved in loom file format using Velocyto package (*84*). The loom files from single samples were combined using scVelo package (*85*)(V 0.1.16) and were normalized with default parameters using “scVelo.pp.filter_and_normalize”. First and second-order moments were calculated for each cell across its 30 nearest neighbors in PCA space (scVelo.pp.moments(n_pcs=15, n_neighbors=30). Velocity was estimated and the velocity graph was constructed using default parameters in “stochastic” mode (scv.tl.velocity(mode=‘stochastic’);scv.tl.velocity_graph). PAGA was used to quantify the transition between clusters (partitions) using PAGA algorithm as implemented in scVelo with the default parameters (scv.tl.paga(groups=‘clusters’,use_rna_velocity=True)). Transitions usually ranged between 0 and 1, and a higher value means a better transition confidence. For visualization we multiplied these values by 100. All transitions are shown in Figure 4A, and those passing a transition confidence threshold (>10; default=5) for nodes from or to NSCs are in Figure 4B.

#### Human Normotypic Brain Specimens

Human normotypic rapid post-mortem brain tissue samples were obtained from the Neuroimmunology Research Laboratory, Centre de Recherche du Centre Hospitalier de l’Université de Montréal under ethics approval protocol BH07.001. Brains were cut in the coronal plane and immersed in 3% paraformaldehyde or formalin for 1–21iweeks and then portions of their lateral ventricular walls were excised.

#### Immunohistochemistry

Slides with 5μm thick sections were baked overnight at 60°C then de-paraffinized and rehydrated using a graded series of xylene and ethanol, respectively. For heat-mediated antigen retrieval, slides were incubated in citrate buffer at 125°C for 20 minutes in a decloaking chamber (BioCare Medical), followed by a cool-down period. Slides were rinsed in distilled water and phosphate buffered saline (PBS) and blocked using Protein Block (Spring Bioscience) for 30 minutes. Sections were incubated with primary antibodies diluted in 2% BSA in PBS overnight in a humid chamber at 4°C, washed using the immunofluorescence (IF) buffer (0.05% Tween-20 and 0.2% Triton X-100 in PBS) and incubated with secondary antibodies diluted in 2% BSA in PBS for 1 hour at room temperature. Following additional wash steps in IF buffer, the slides were mounted with ProLong™ Diamond Antifade Mountant with DAPI (Invitrogen) to counterstain cell nuclei. Fluorescent images were acquired using a ZEISS LSM 700 laser scanning confocal microscope.

#### Antibodies

The following antibodies were used for immunolabelling: VIM (BL202 – EMD Millipore); SOX9 (ab185966 -Abcam); GFAP (ab4674 – Abcam); OLIG2 (ab9610 – EMD Millipore); ASCL1 (ab74065 – Abcam); TMEM119 (E3E4T – Cell Signaling); EGFR (ab32198 – Abcam), CLDN5 (LS-C352946 – LS Bio); ANXA1 (ab214486 – Abcam); MBP (ab40390 – Abcam); TAGLN2 (ab121146 – Abcam); CD3D (MA5-32462 – ThermoFisher).

#### Pathway and Biological Process Enrichment Analysis

Pathway and biological process enrichment analysis was performed using Metascape (multi-list enrichment analysis) (86). For each gene list a maximum of the top 500 differentially expressed genes were used for each cell type. Pathway and biological process enrichment analysis was carried out with the following ontology sources: KEGG Pathway, GO Biological Processes, Reactome Gene Sets, Canonical Pathways, and WikiPathways. All genes in the genome were used as the enrichment background. Terms with a p-value < 0.01, a minimum count of 3, and an enrichment factor > 1.5 were collected. p-values were calculated based on the cumulative hypergeometric distribution, and q-values were calculated using the Benjamini-Hochberg procedure to account for multiple tests.

## ACKNOWLEDGEMENTS

We would like to thank Carmen Sabau for her contribution to the administrative work of the project, as well as Rozica Bolovan for technical support.

## FUNDING

Cancer Research Society 70813 (KP)

Canadian Cancer Research Institute 702411 (KP)

Brain Tumour Foundation of Canada 62766 (KP

TARGiT Foundation (KP)

A Brilliant Night Foundation (KP)

Argento Family Group Ercole (KP).

KP is supported by a clinician-scientist salary award from the Fonds de Recherche du Québec and the William Feindel Chair in Neuro-Oncology.

## Author Contributions

Conceptualization: SB, KP

Methodology: SB, JN, RA, KP

Investigation: SB, JN, RA, PL, ML, AD, MS, AP, JA, MCG, KP

Visualization: SB, JN, PL, AD, KP

Funding acquisition: KP

Project administration: KP

Supervision: KP

Writing – original draft: SB, JN, PL, KP

Writing – review & editing: SB, JN, PL, ML, KP

## Data and Materials Availability

All data are available in the main text or the supplementary materials.

