## Supplementary material for "Identity and Nature of Neural Stem Cells in the Adult Human Subventricular Zone": Figure S1

| SAMPLE | AGE | SEX | DIAGNOSIS |
|---|---|---|---|
| SVZ337 | 53 | M | Astrocytoma grade 3 |
| SVZ350 | 51 | M | Glioblastoma |
| SVZ359 | 51 | M | Astrocytoma grade 3 |
| SVZ373 | 62 | M | Glioblastoma |
| SVZ375 | 70 | F | Metastatic adenocarcinoma |
| SVZ376 | 55 | M | Glioblastoma |
| SVZ379 | 38 | M | Astrocytoma grade 2 |
| SVZ383 | 63 | M | Glioblastoma |
| SVZ391 | 62 | M | Large cell carcinoma |
| SVZ395 | 72 | F | Glioblastoma |
| SVZ400 | 72 | F | Glioblastoma |
| SVZ420 | 57 | F | Glioblastoma |
| SVZ423 | 64 | F | Glioblastoma |
| SVZ428 | 45 | M | Glioblastoma |
| SVZ444 | 71 | F | Glioblastoma |

**Figure S1. Subventricular zone sample acquisition demographics.**
