## Supplementary figures and images for "Identity and Nature of Neural Stem Cells in the Adult Human Subventricular Zone"

### Figure S2

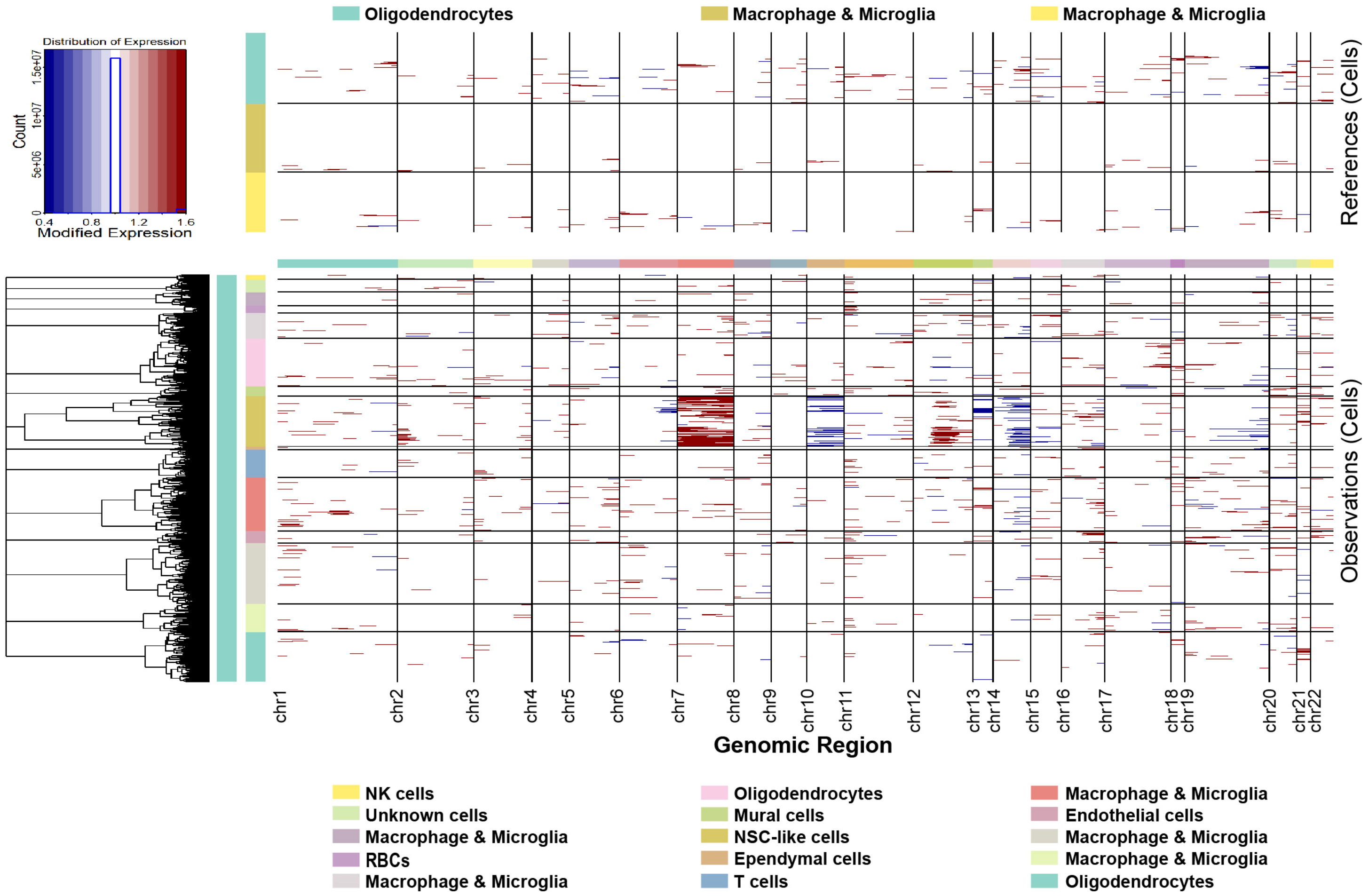


**Figure S2. Copy number alteration analysis of all subventricular cells.**
